# Parametric information about eye movements is sent to the ears

**DOI:** 10.1101/2022.11.27.518089

**Authors:** Stephanie N Lovich, Cynthia D King, David LK Murphy, Rachel Landrum, Christopher A Shera, Jennifer M Groh

**Author notes:** These authors contributed equally to this work and are listed in reverse chronological order of their period of most intense involvement (Stephanie N Lovich, Cynthia D King, David LK Murphy).

## Abstract

Eye movements alter the relationship between the visual and auditory spatial scenes. Signals related to eye movements affect neural pathways from the ear through auditory cortex and beyond, but how these signals contribute to computing the locations of sounds with respect to the visual scene is poorly understood. Here, we evaluated the information contained in eye movement-related eardrum oscillations (EMREOs), pressure changes recorded in the ear canal that occur in conjunction with simultaneous eye movements. We show that EMREOs contain parametric information about horizontal and vertical eye displacement as well as initial/final eye position with respect to the head. The parametric information in the horizontal and vertical directions can be modelled as combining linearly, allowing accurate prediction of the EMREOs associated with oblique (diagonal) eye movements. Target location can also be inferred from the EMREO signals recorded during eye movements to those targets. We hypothesize that the (currently unknown) mechanism underlying EMREOs could impose a two-dimensional eye-movement related transfer function on any incoming sound, permitting subsequent processing stages to compute the positions of sounds in relation to the visual scene.

**Significance Statement:** When the eyes move, the alignment between the visual and auditory scenes changes. We are not perceptually aware of these shifts -- which indicates that the brain must incorporate accurate information about eye movements into auditory and visual processing. Here we show that the small sounds generated within the ear by the brain contain accurate information about contemporaneous eye movements in the spatial domain: the direction and amplitude of the eye movements could be inferred from these small sounds. The underlying mechanism(s) likely involve(s) the ear’s various motor structures, and could facilitate the translation of incoming auditory signals into a frame of reference anchored to the direction of the eyes and hence the visual scene.

## Introduction

Every time we move our eyes to localize multisensory stimuli, our retinae move in relation to our ears. These movements shift the alignment of the visual scene (as detected by the retinal surface) with respect to the auditory scene (as detected based on timing, intensity, and frequency differences in relation to the head and ears). Precise information about each eye movement is therefore needed to connect the brain’s views of visual and auditory space to one another (e.g. e.g. 1, 2, 3). Most previous work about how eye movement information is incorporated into auditory processing has focused on cortical and subcortical brain structures (4–24), but the recent discovery of eye-movement related eardrum oscillations (EMREOs) (25–28) suggests that the process might manifest much earlier in the auditory periphery. EMREOs can be thought of as a biomarker of underlying efferent information impacting the internal structures of the ear in association with eye movements. What information this efferent signal contains is currently uncertain.

We reasoned that if this efferent signal is to play a role in linking auditory and visual space across eye movements, EMREOs should be *parametrically* related to the associated eye movement. Specifically, EMREOs should vary in a regular and predictable fashion with both horizontal and vertical displacements of the eyes, and some form of information regarding the initial position of the eyes should also be present. These properties are required if the efferent signal underlying EMREOs is to play a role in linking hearing and vision. Notably, this parametric relationship is not required of alternative possible roles, such as synchronizing visual and auditory processing in time or enhanced attentional processing of sounds regardless of their spatial location (29–33).

Accordingly, we evaluated the parametric spatial properties of EMREOs in human participants by varying the starting and ending positions of visually-guided saccades in two dimensions. We find that EMREOs do in fact vary parametrically depending on the saccade parameters in both horizontal and vertical dimensions and as a function of both initial eye position in the orbits and the change in eye position relative to that initial position. EMREOs associated with oblique (diagonal) saccades can be predicted by the linear combination of the EMREOs associated with strictly horizontal and vertical saccades. Furthermore, an estimate of target location can be decoded from EMREOs alone – i.e. where subjects looked in space can be roughly determined from their observed EMREOs.

These findings suggest that the eye-movement information needed to accomplish a coordinate transformation of incoming sounds into a visual reference frame is fully available in the most peripheral part of the auditory system. While the precise mechanism that creates EMREOs remains unknown, we propose that the underlying mechanisms might introduce a transfer function to the sound transduction process that serves to adjust the gain, latency, and/or spectral dependence of responses in the cochlea. In principle, this could provide later stages of auditory processing access to an eye-centered signal of sound location for registration with the eye-centered visual scene (1), Indeed, recent work has shown that changes in muscular tension on the ossicular chain would be expected to affect gain and latency of sound transmission through the middle ear, thus supporting the plausibility of this hypothesis (34, 35).

## Results

We used earbud microphones to record internally-generated oscillations in the ear canals of human subjects with normal hearing and corrected to normal vision. All procedures concerning human participants were approved by the Duke University Institutional Review Board (see Supplementary Methods), and all participants provided informed consent before beginning the experiments.

Participants performed eye-movement tasks involving various visual fixation and target configurations (Figure 1 inset; Supplementary Figure 1). No external sounds were presented in any task. At the beginning of each trial, subjects fixated a visual fixation point for a minimum of 200 ms and then made a saccade to a second target, which they then fixated for another 200 ms (Supplementary Figure 1a). Any trials with micro- or corrective saccades during the 200 ms prior to or following main fixation-point-to-target saccade were discarded, to ensure a stable baseline ear-canal recording could be established without intrusions by other eye movements.

**Figure 1.**
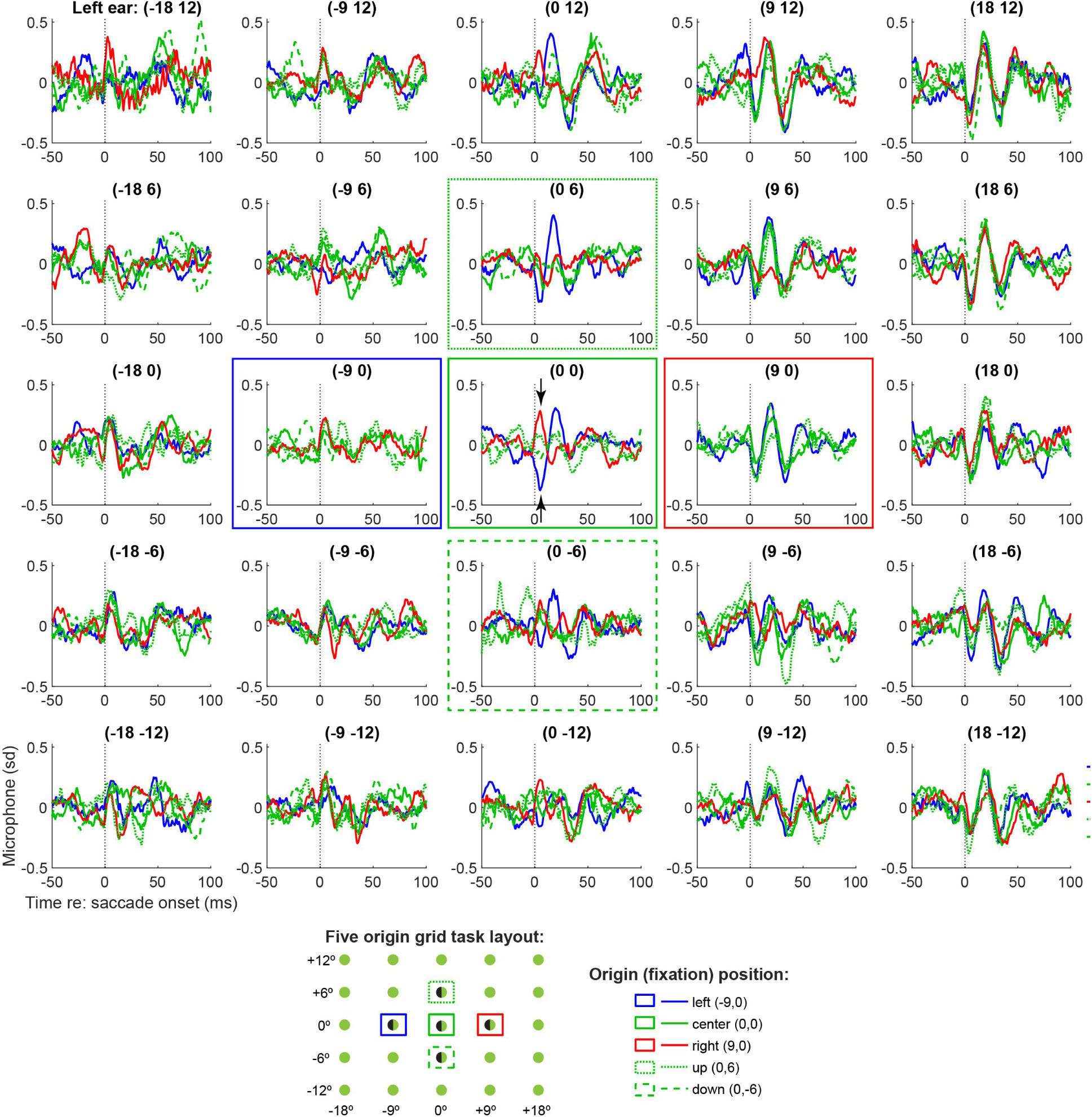
EMREOs recorded during the five-origin grid task. Each panel shows the grand average EMREO signal generated when saccades were made to that location on the screen (average of N=10 subjects’ individual left ear averages). For example, the top right panel shows microphone recordings during saccades to the top right (contralateral) target location, and the color and line styles of each trace in that panel correspond to saccades from different initial fixation points. e.g. the red traces originated from the rightward fixation, the blue from the leftward fixation etc as indicated by the legend and boxes of the same color and line style. Both magnitude and phase vary as a function of initial eye position and target location, with contralateral responses being larger than ipsilateral. Phase reversal occurs based on the location of the target with respect to the initial fixation position, as can be seen for the central target location (central panel), where the EMREOs evoked for saccades from the rightward fixation (red traces) show an opposite phase relationship as those evoked for saccades from the leftward fixation (blue traces). Corresponding grand averages for right ear data are shown in Supplementary Figure 3. These data are presented in Z-units; the peak-equivalent sound levels for 18 degree horizontal targets are roughly 55-56 dB SPL; see Supplementary Figure 2 for the mean and distributions across the subject population (range ∼49-64 dB SPL).

We first tested subjects (N=10) on a task involving variation in both initial fixation position and target location along both horizontal and vertical dimensions – the “five-origin grid task”. Subjects fixated on an initial fixation light located either straight ahead, 9° left or right, or 6° up or down, and then made a saccade to a target located within an array of possible target locations spanning +/- 18° horizontally and +/- 12° vertically (Figure 1 inset; Supplementary Figure 1b). Results of this task are shown in Figure 1. Each panel shows the average microphone signal recorded in the left ear canal (averaged across all subjects) associated with saccades to a target at that location – e.g. the top right panel shows all saccades to the top right target location. The color and line styles of the waveforms correspond to the five initial fixation positions from which the saccades could originate in space.

The first overall observation from this figure is that the *magnitude* of the waveform of the EMREO depends on both the *horizontal* and *vertical* dimensions. In the horizontal dimension, EMREOs are larger for more contralateral target locations: compare the column on the right (contralateral) to the column on the left (ipsilateral). The pattern is reversed for right ear canal recordings (Supplementary Figure 3). In the vertical dimension, EMREOs are larger for higher vs lower targets in both left and right ears (compare top row to bottom row in Figure 1/Supplementary Figure 3).

The second overall observation from this figure is that the *phase* of the EMREO waveform depends on the horizontal location of the target *with respect to* the fixation position. Specifically, the first deflection after saccade onset is a peak for the most ipsilateral targets (left-most column) and trough for the most contralateral targets (right-most column). But *where* this pattern reverses depends on the initial fixation position. Specifically, consider the red vs blue traces in the middle column of the figure, which correspond to targets along the vertical meridian. Red traces involve saccades to these targets from the fixation position on the right, and thus involve leftward (ipsiversive) saccades. The red traces in this column begin with a peak followed by a trough. In contrast, the blue traces involve saccades to these targets from the fixation position on the left, i.e. rightward or contraversive saccades. The blue traces begin with a trough followed by a peak. The pattern is particularly evident in the central panel (see arrows).

The phase reversal as a function of the combination of target location and initial eye position suggests that the EMREO waveforms might align *better* when plotted in an eye-centered frame of reference. Figure 2 demonstrates that this is indeed the case: the data from Figure 1 is re-plotted as a function of target location *relative* to the initial fixation position. The eight panels around the center represent the traces for the subset of targets that can be fully analyzed in an eye-centered frame, i.e. the targets immediately left, right, up, down, and diagonal relative to the five fixation locations. By plotting the data based on the relative location of the targets to the origins, the waveforms are better aligned, showing no obvious phase reversals.

**Figure 2.**
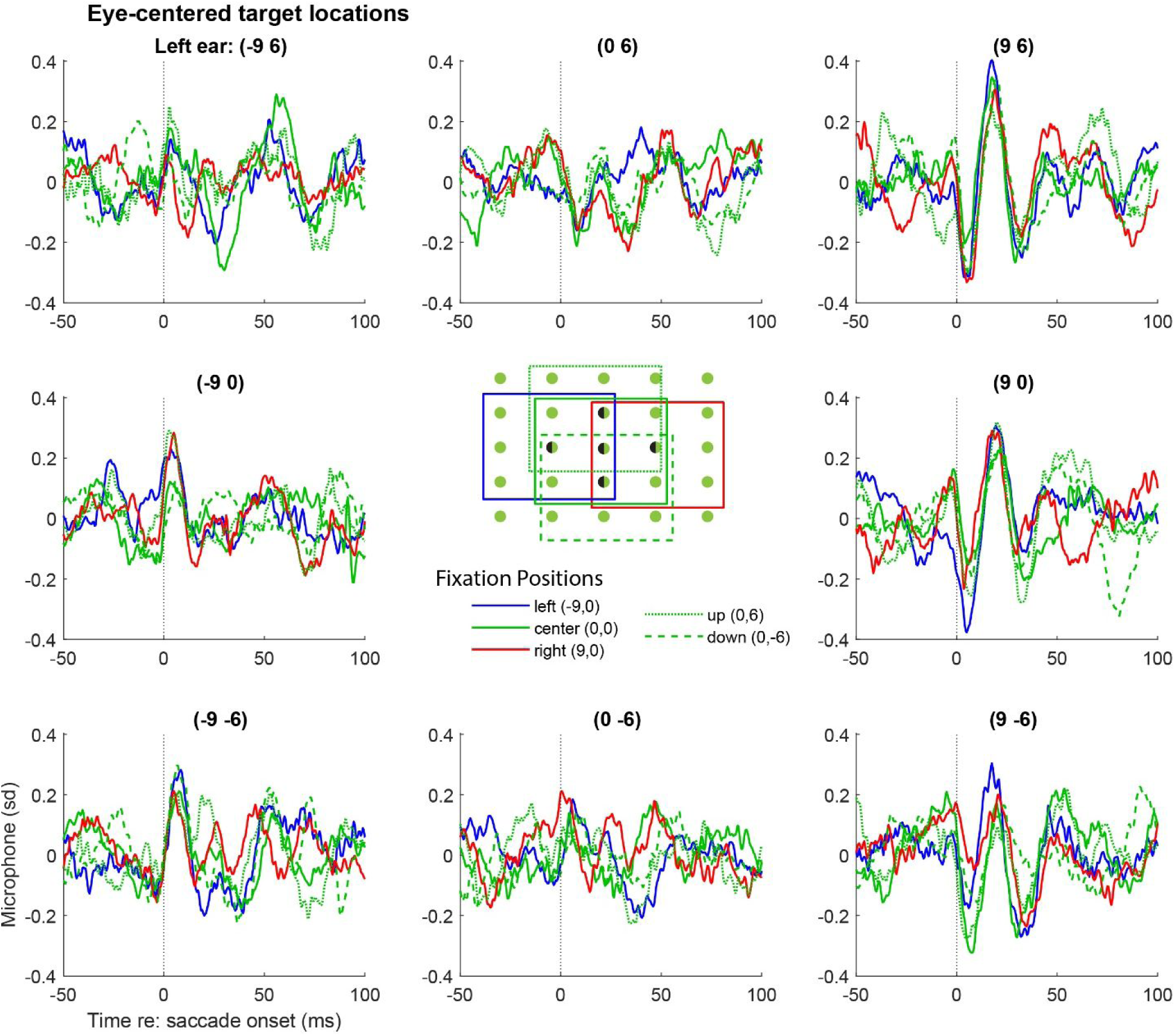
Replotting the grand average EMREOs as a function of relative target location shows better, but not perfect, correspondence of the EMREOs across different fixation positions. The data shown are a subset of those shown in Figure 1, but here each panel location corresponds to a particular target location defined <u>relative</u> to the associated fixation position. The color/linestyle indicates the associated relative fixation position. For example, the waveforms in the upper right panel all involved 9° rightward and 6° upward saccades; the red trace in that panel indicates those that originated from the 9° right fixation; the blue those from the 9° left fixation etc. Only relative target locations that existed for all 5 fixation positions are plotted, as indicated by the inset. Corresponding right ear data are shown in Supplementary Figure 4.

Although the waveforms are better aligned when plotted based on target location *relative* to initial eye position, some variation related to that fixation position is still evident in the traces. That is, in each panel, the EMREO waveforms with different colors/line styles (corresponding to different fixation positions) do not necessarily superimpose perfectly. This suggests that a model that incorporates both relative target position and original fixation position, in both horizontal and vertical dimensions, is needed to account for the findings. Furthermore, a statistical accounting of these effects is needed. Accordingly, we fit the data to the following regression equation:

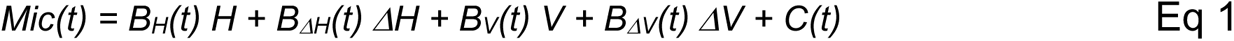

where *H* and *V* correspond to the initial horizontal and vertical eye position and *ΔH* and *ΔV* correspond to the respective changes in position associated with that trial. The slope coefficients *B_H_, B_ΔH_, B_V_, B_ΔV_*are time-varying and reflect the dependence of the microphone signal on the respective eye position/movement parameters. The term *C(t)* contributes a time-varying “constant” independent of eye movement metric and can be thought of as the best fitting average oscillation across all initial eye positions and changes in eye position. We used the measured values of eye position/change in eye position for this analysis rather than the associated fixation and target locations so as to incorporate trial-by-trial variability in fixation and saccade accuracy.

This model is a conservative one, assessing whether a linear relationship between the microphone signal and the relevant eye position/movement variables can provide a satisfactory fit to the data. As such, it provides a lower bound but does not preclude that higher quality fits could be achieved via non-linear modeling. This approach is similar to the general linear models applied to fMRI data (e.g. (36) and differs chiefly in that we make no assumptions about the underlying temporal profile of the signal (such as a hemodynamic response function) but allow the temporal pattern to emerge in the time-varying fits of the coefficients.

Figure 3 shows the average of these time-varying values of the slope coefficients across subjects (blue = left ear; red = right ear) and provides information about the contribution of these various eye movement parameters to the EMREO signal. A strong, consistent dependence on horizontal eye displacement is observed, consistent with our previous report (Figure 3a) (25). This component is oscillatory and begins slightly before the onset of the eye movement, inverting in phase for left vs right ears. The thickened parts of the line indicate periods of time when this coefficient differed significantly from 0 with 95% confidence (Shaded areas are +/-SEM). There is also an oscillatory and binaurally phase-inverting signal related to the initial position of the eyes in the horizontal dimension (Figure 3b). This signal is smaller and more variable across subjects.

**Figure 3.**
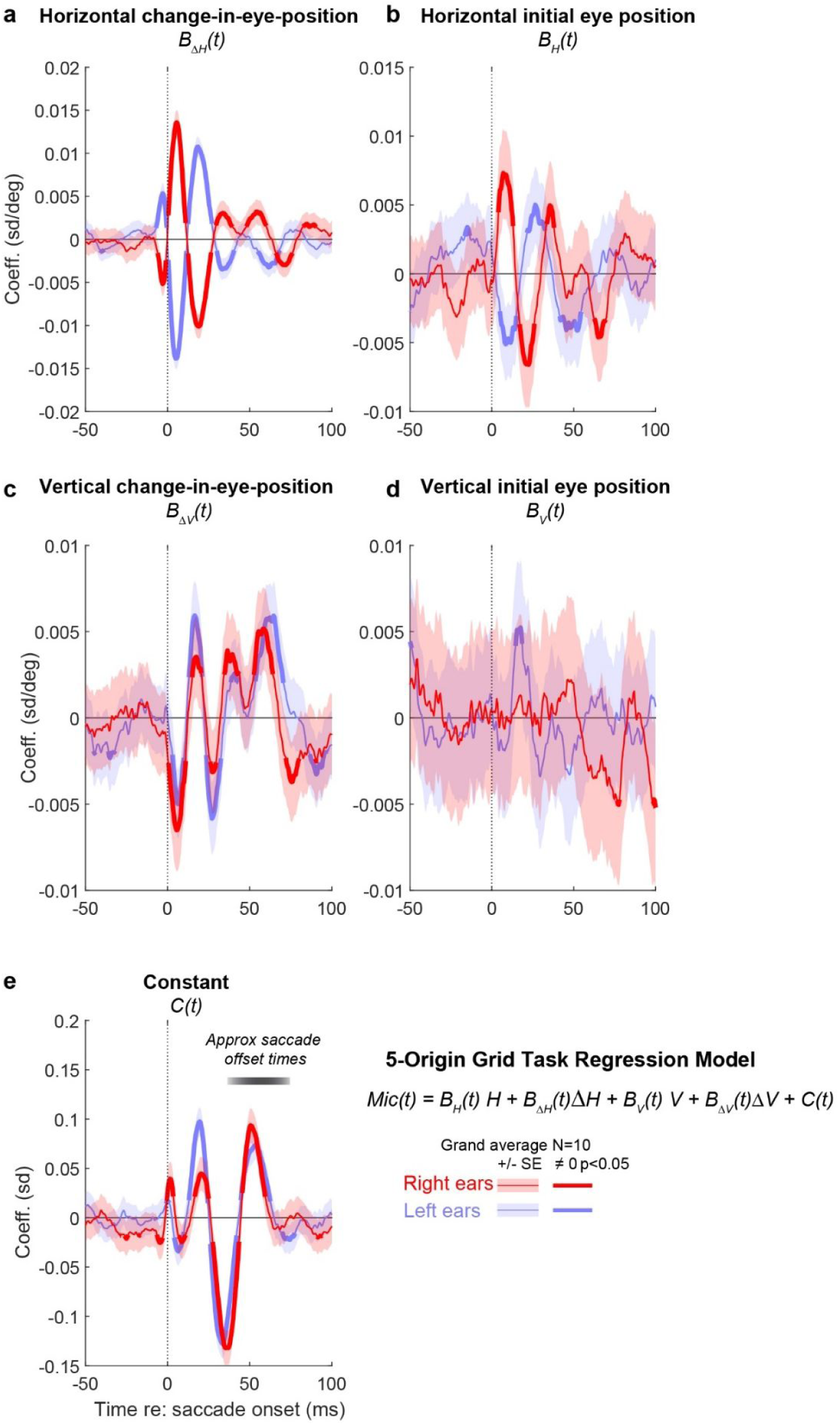
Regression analysis of EMREOs shows contributions from multiple aspects of eye movement: horizontal and vertical change-in-eye-position **(a, c)**, horizontal initial eye position **(b)**, as well as a constant component that was consistent across saccades **(e)**. The contribution of vertical initial eye position was weaker (**d**). The regression involved modeling the microphone signal at each time point, and each panel shows the time varying values of the coefficients associated with the different aspects of the eye movement (horizontal vs. vertical, change-in-position and initial position). The regressions were fit to individual subjects’ microphone recordings, and plotted here as grand averages of these regression coefficients across the N=10 subjects tested in the 5-origin grid task. Microphone signals were z-scored in reference to baseline variability during a period −150 to 120 ms prior to saccade onset. Results are presented in units of standard deviation (panel e) or standard deviation per degree (panels **a-d**). Shaded areas represent +/-SEM.

In the vertical dimension, the effect of vertical saccade amplitude is in phase for both the left and right ears; it exhibits an oscillatory pattern, although not obviously sinusoidal like the one observed for the horizontal saccade amplitude. Initial position of the eyes in the vertical dimension exerts a variable effect across participants such that it is not particularly evident in this grand average analysis; this may be related to poorer abilities to localize sounds in the vertical vs. horizontal dimensions (37–40).

Finally, there is a constant term that is similar in the two ears and peaks later with respect to saccade onset than is the case for the other coefficients (Figure 3e). As noted above, this constant term can be thought of as encapsulating the average EMREO waveform that occurs when pooling across all the eye movements in the dataset, regardless of their initial positions or horizontal or vertical components.

We next investigated whether the fits obtained in one task match those obtained in a different task. We reasoned that if the information contained in the EMREO signal reflects the eye movement itself, then task context should not matter. Furthermore, the regression model should provide a good way to accomplish this comparison since it does not require that the exact same locations and eye movements be tested across tasks.

To test these questions, we collected data using two simplified tasks, the single-origin-grid task (with a single initial fixation in the center, Supplementary Figure 1c) and the horizontal/vertical task (with a single fixation at the center and targets on the horizontal and vertical meridians, generating purely horizontal or vertical saccades, Supplementary Figure 1d). Ten subjects (four of whom also completed the 5-origin grid task) completed both the single-origin grid task and the horizontal/vertical saccade. We fit the results from these tasks using the same regression procedure but omitting the initial fixation position terms, i.e.:

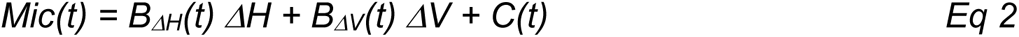

As shown in Figure 4, both tasks yield similar values of the regression coefficients for horizontal change-in-position (*B_ΔH_(t)*) and the constant term (*C(t)*) (grand average across the population, black vs. green traces). The vertical change-in-position term (*B_ΔV_(t)*) was slightly more variable but also quite consistent across tasks.

**Figure 4.**
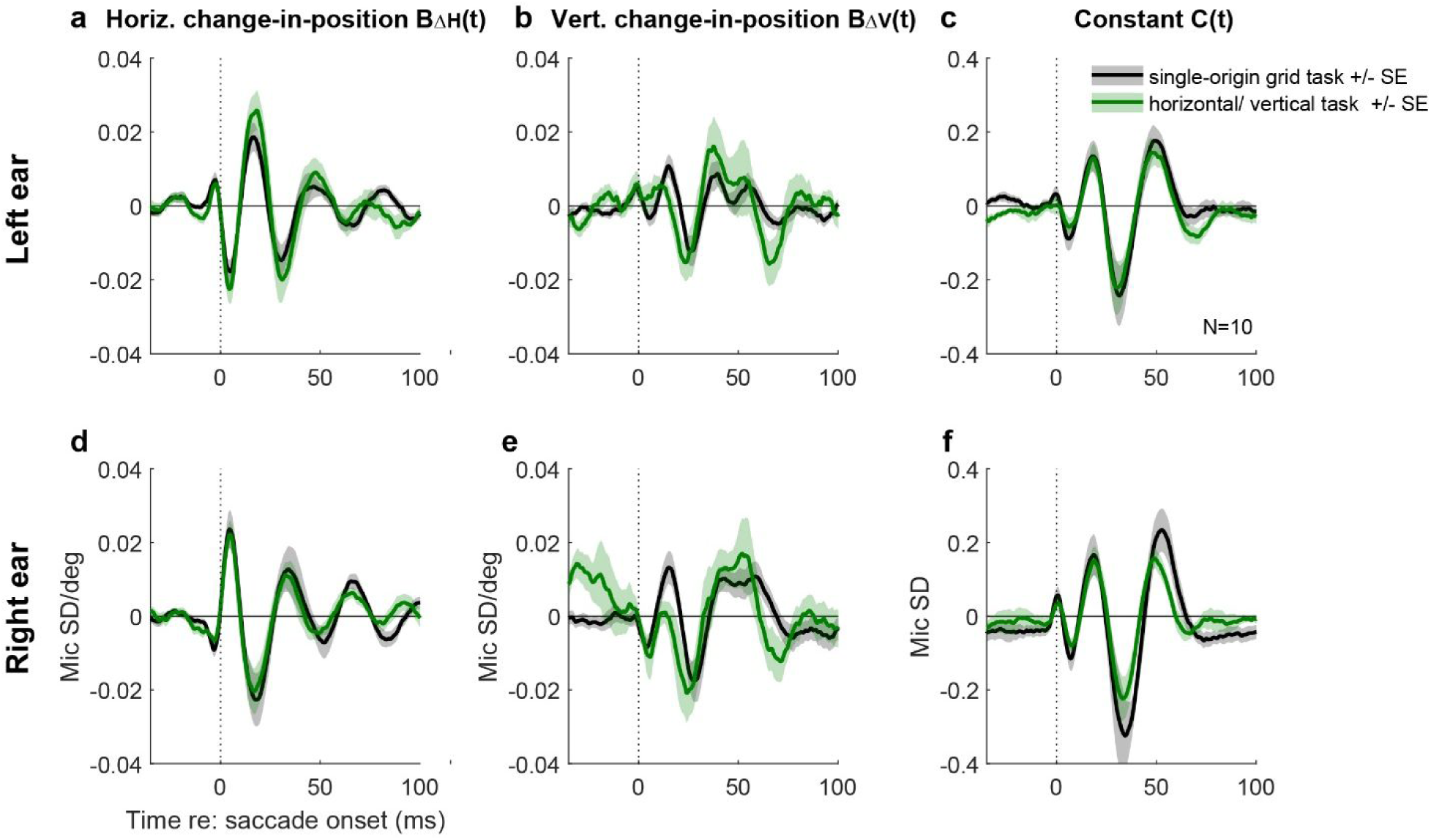
Different tasks generate similar regression coefficient curves. Grand average of the regression results for the single-origin grid (black lines) and horizontal/vertical (green lines) tasks. The lines and shading represent the average and standard error of the coefficient values across the same 10 subjects for the two tasks. See Supplementary Figure 5 for corresponding findings among the individual subjects.

Given the consistency of the regression coefficient values between the single-origin grid and horizontal/vertical tasks (and see (26) for similar results involving spontaneous vs. task-guided eye movements), we surmised that it should be possible to use the *coefficient* values from one task to predict the EMREO *waveforms* in the other. Specifically, we used the time-varying regression values from purely horizontal and purely vertical saccades in the horizontal/vertical task to predict the observed waveforms from oblique saccades in the single origin grid task. This method can be used to evaluate the quality of the regression-based EMREO prediction not only for target locations tested in both tasks, i.e. the horizontal and vertical meridians, but also for oblique targets tested only in the grid task.

The black traces in Figure 5 show the grand average microphone signals associated with each target in the single-origin grid task. The location of each trace corresponds to the physical location of the associated target in the grid task (similar to Figure 1). The superimposed predicted wave forms (red traces) were generated from the B_ΔH_(t), B_ΔV_(t), and C(t) regression coefficients fit to only the horizontal/vertical data, then evaluated at each target location and moment in time to produce predicted curves for each of the locations tested in the grid task.

**Figure 5.**
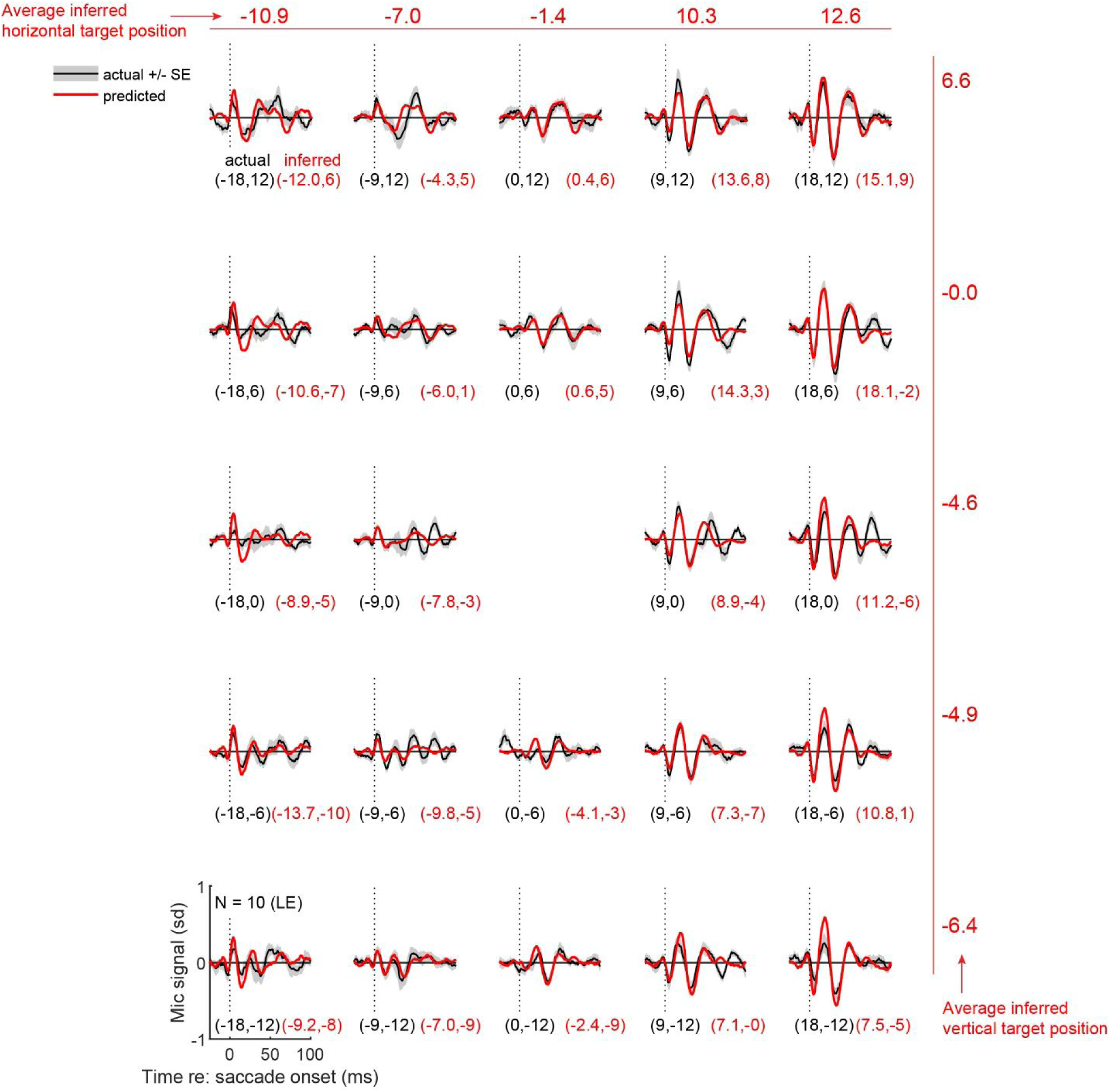
Regression coefficients fit to microphone recordings from the horizontal/vertical-saccade task can be used to predict the waveforms observed in the grid task and their corresponding target locations. Combined results for all N=10 participants’ left ears. The black traces indicate the grand average of all the individual participants’ mean microphone signals during the single-origin grid task, with the shading indicating +/- the standard error across participants. The red traces show an estimate of the EMREO at that target location based only on regression coefficients measured from the horizontal/vertical task. Black values in parentheses are the actual horizontal and vertical coordinates for each target in the grid task. Corresponding red values indicate the inferred target location based on solving a multivariate regression which fits the observed grid task microphone signals in a time window (−5 to 70 ms with respect to saccade onset) to the observed regression weights from the horizontal/vertical task for each target location. The averages of these values in the horizontal and vertical dimensions are shown across the top and right sides. See Figure 6 for additional plots of the inferred vs actual target values, and Supplementary Figure 6 for corresponding right-ear data.

Overall, there is good correspondence between the predicted EMREO oscillations and the observed EMREO from actual microphone recordings, including the oblique target locations that were not tested in the horizontal/vertical task. This illustrates two things: 1) the EMREO is reproducible across task contexts, and 2) the horizontal and vertical change-in-position contributions interact in a reasonably independent way, so that the EMREO signal observed for a combined horizontal-vertical saccade can be predicted as the sum of the signals observed for purely horizontal and purely vertical saccades with the corresponding component amplitudes.

Given that it is possible to predict the microphone signal from one task context to another, it should also be possible to decode the target location associated with an eye movement from just the simultaneously-recorded microphone signal. To do this, we again used the weights from the horizontal/vertical task data for the regression equation:

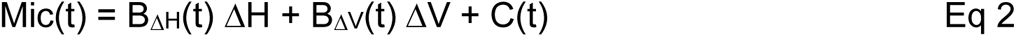

Specifically, we used the Mic(t) values observed in the single-origin grid task to solve this system of multivariate linear equations across the time window −5 to 70 ms with respect to the saccade (a time period in which the EMREO appears particularly consistent and substantial in magnitude) to generate the “read out” values of ΔH and ΔV associated with each target’s actual ΔH and ΔV. We conducted this analysis on the left ear and right ear data separately. The left ear results of this analysis are seen in each of the individual panels of Figure 5; the black values (e.g. −18, 12) indicate the actual horizontal and vertical locations of the target, and the associated red values indicate the inferred location of the target. Across the top of the figure, the numbers indicate the average inferred horizontal location, and down the right side, the numbers indicate the average inferred vertical location. These results indicate that, on average, the targets can be read out in the proper order, but the spatial scale is compressed: the average read-out values for the +/-18 degree horizontal targets are +/- ∼11-12 degrees, and the averages for the vertical +/- 12 degree targets are +/- ∼6-7 degrees. Similar patterns occurred for the right ear data (Supplementary Figure 6).

Plots of these target readouts in both horizontal and vertical dimensions for both ears are shown in Figure 6a-f. Figure 6a shows the inferred location of the target (red dots) connected to the actual location of the target (black dots) using the data from Figure 5, i.e the left ear readout, and Figure 6b-c show regressions of these target readouts as a function of the horizontal and vertical locations. Figure 6d-f show the corresponding results for the right ears. Altogether, these figures illustrate that the readout accuracy is better in the horizontal than in the vertical dimensions. Quantitatively, the r^2^ values for the horizontal dimension were 0.89 (LE) and 0.91 (RE), and the corresponding values for the vertical dimension were 0.61 (LE) and 0.67 (RE). Slopes were also closer to a value of 1 (the ideal) for the horizontal dimension (0.71, LE; 0.77, RE) than for the vertical dimension (0.51, LE, 0.51, RE).

**Figure 6.**
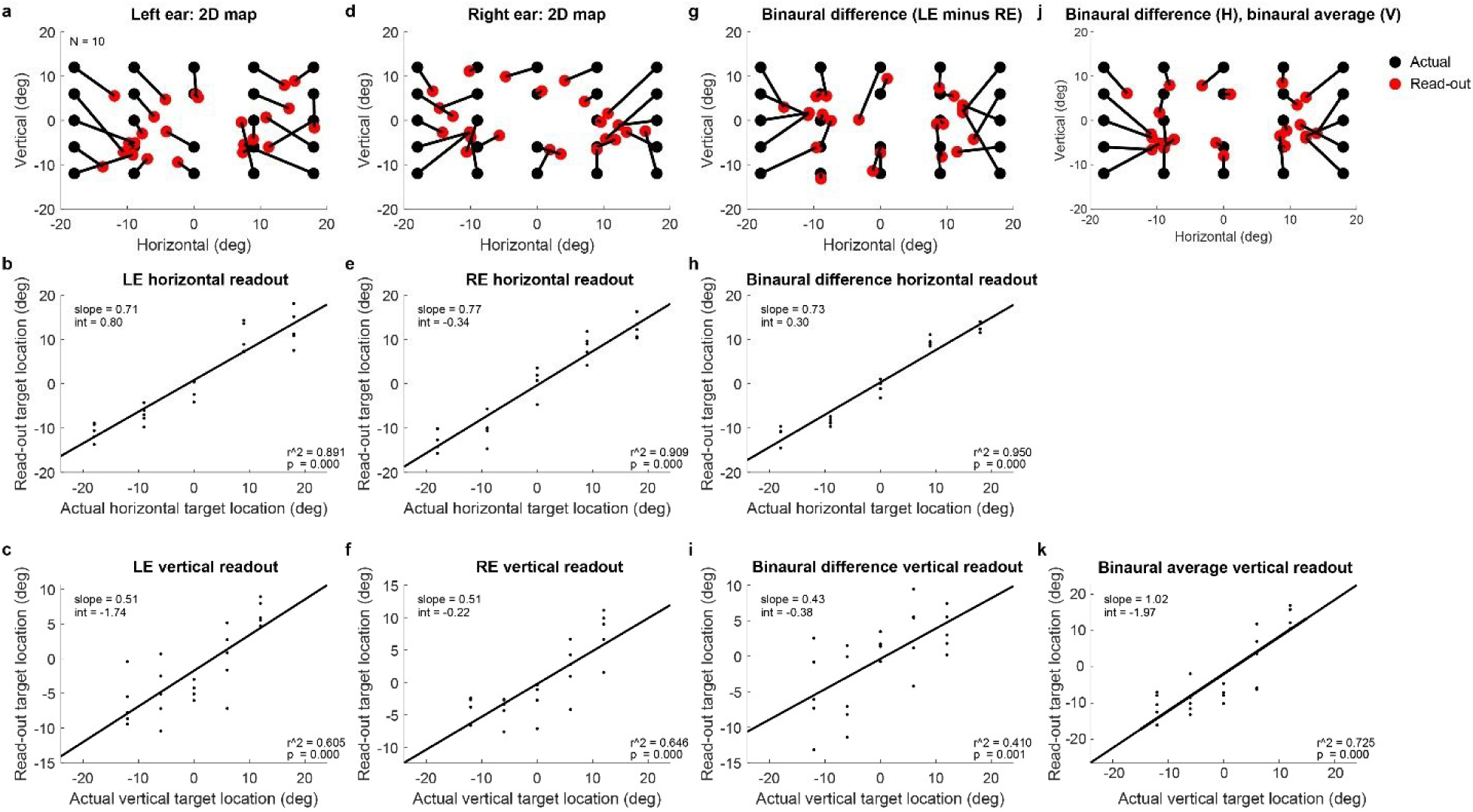
Multiple ways of reading out target location from the ear canal recordings. As in Figure 5 and Supplementary Figure 6, the relationship between EMREOs and eye movements was quantitatively modelled using Eq 2 and the ear canal data recorded in the horizontal/vertical task. Inferred grid task target location was “read out” by solving equation (2) for ΔH and ΔV using the coefficients as fit from the horizontal/vertical task and the microphone values as observed in the single-origin grid task; see main text for details. **a**. Inferred target location (red) compared to actual target location (black), based on the left ear (same data as in Figure 5). **b**. Horizontal component of the read-out target vs the actual horizontal component (left ear microphone signals). **c**. Same as **(b)** but for the vertical component. **d-f.** Same as A-C but for the right ear. **g-i**, Same as a-c and d-f but computed using the binaural difference between the microphone signals (left ear – right ear). **j., k.** A hybrid read-out model **(j**) using binaural difference in the horizontal dimension **(h)** and binaural average in the vertical dimension **(k)**. Related findings at the individual subject level are provided in Supplementary Figure 7.

Given that it is known that the brain uses binaural computations for reconstructing auditory space, we wondered whether the accuracy of this read-out could be improved by combining signals recorded in each ear simultaneously. We first considered a binaural difference computation, subtracting the right ear microphone recordings from the left, thus eliminating the part of the signal that is common between the two ears. Figure 6g shows the results. Generally, the horizontal dimension is well ordered whereas the vertical dimension continues to show considerable shuffling. This can also be seen in Figure 6h and 6i, which show the relationship between the inferred target location and the true target location, plotted on the horizontal and vertical dimension, respectively. The correlation between inferred and actual target is higher in the horizontal dimension (r^2^ 0.95) than the vertical dimension (r^2^ 0.41), which is actually worse than the monaural readouts. This makes sense because the binaural difference computation serves to diminish the contribution from aspects of the signal that are in phase across the two ears, such as the dependence on vertical change in eye position. We then reasoned that improvement in the vertical readout could be achieved by instead averaging, rather than subtracting, the signals across the two ears, and indeed this is so: averaging across the two ears produces an improved vertical readout (r^2^ 0.73, Figure 6k). Finally, a hybrid readout operation in which the horizontal location is computed from the binaural difference, and the vertical location is computed from the binaural average, produces an additional modest improvement, yielding the best overall reconstruction of target location (Figure 6j).

We next considered how well this readout operation performed at the level of individual subjects. Results for each subject are shown in Supplementary Figure 7 a-j, and a population summary is shown in Supplementary Figure 7k. The relationship between the inferred location and the actual location was statistically significant (p<0.05) for all 10 subjects in the horizontal dimension, and for 7 of 10 subjects in the vertical dimension. This confirms that we can predict the horizontal location of the target of a saccade from ear recordings in each individual subject, and the vertical location can be predicted for most, but not all, subjects.

Finally, we evaluated the error when reading out the target location of individual trials. The preceding analyses show the results when the read-out operation is performed on the average waveform observed across trials for a given target location. The same readout can also be computed for each individual trial. When conducted on all individual trials, the resulting scatter can be computed as the average standard deviation observed across target locations and subjects. We found that the average scatter or standard deviation was 19.1 degrees in the horizontal dimension and 23.8 degrees in the vertical dimension. For the horizontal dimension, this corresponds roughly half of the range of space tested (+/- 18 degrees), whereas in the vertical dimension this corresponds to nearly the whole range (+/- 12 degrees).

Overall, these results parallel human sound localization which relies on a binaural difference computation in the horizontal dimension (and is more accurate in that dimension), vs. potentially monaural or averaged spectral cues for the vertical dimension (which is less accurate) (41, 42). Indeed, horizontal and vertical sound localization show different patterns of dependence on the loudness of the target sound relative to background noise, further supporting that these operations are accomplished via distinct mechanisms (43).

## Discussion

Sound locations are inferred from head-centered differences in sound arrival time, intensity, and spectral content, but visual stimulus locations are inferred from eye-centered retinal locations (41, 42). Information about eye movements with respect to the head/ears is critical for connecting the visual and auditory scenes to one another (1). This insight has motivated a number of previous neurophysiological studies in various brain areas in monkeys and cats, all of which showed that changes in eye position affected the auditory response properties of at least some neurons in the brain area studied (Inferior colliculus: (8–12); auditory cortex: (5–7); superior colliculus: (18–24); frontal eye fields: (13, 44); intraparietal cortex: (14–17)).

These findings raised the question of where signals related to eye movements first appear in the auditory processing stream. The discovery of EMREOs (25–28, 45) introduced the intriguing possibility that the computational process leading to visual-auditory integration might be manifested in the most peripheral part of the auditory system. Here we show that the signals present in the ear exhibit the properties necessary for playing a role in this process: these signals carry information about the horizontal and vertical components of eye movements, and display signatures related to both change-in-eye-position and the absolute position of the eyes in the orbits. Because of the parametric information present in the EMREO signal, we were able to predict EMREOs in one task from the EMREOs recorded in another, and even predict the target of eye movements from the simultaneous EMREO recording. These predictions were accomplished using strictly linear methods, a conservative approach providing a lower bound on what can be deduced from these signals. Improvements in the “readout” may come from exploration of more powerful non-linear techniques and/or other refinements such as tailoring the time window used for the readout (here, a generous −5 to 70 ms) or stricter criteria for the exclusion of trials contaminated by noise (see “*Methods: Trial exclusion criteria*”). It should be noted that this read-out presumes knowledge of when the saccade starts and that performance would be substantially poorer if conducted in a continuous fashion across time.

Our present observations raise two key questions: what causes EMREOs and how do those mechanisms impact hearing/auditory processing? The proximate cause of EMREOs is likely to be one or more of the known types of motor elements in the ear^1^: the middle ear muscles (stapedius and tensor tympani), which modulate the motion of the ossicles (46–48), and the outer hair cells, which modulate the motion of the basilar membrane (49). One or more of these elements may be driven by descending brain signals originating from within the oculomotor pathway and entering the auditory pathway somewhere along the descending stream that ultimately reaches the ear via the 5^th^ (tensor tympani), 7^th^ (stapedius muscle), and/or 8^th^ nerves (outer hair cells) (see refs: 50, 51-55) for reviews). Efforts are currently underway in our laboratory to identify the specific EMREO generators/modulators (56, 57) (58).

Uncovering the underlying mechanism should shed light on another question. Does the temporal pattern of the observed EMREO signal reflect the time course and nature of that underlying mechanism’s impact on auditory processing? It is not clear how an oscillatory signal like the one observed here might contribute to hearing. However, it is also not clear that the underlying mechanism is, in fact, oscillatory. Microphones can only detect signals with oscillatory energy in the range of sensitivity of the microphone. It is possible that the observed oscillations reflect ringing associated with a change in some mechanical property of the transduction system, and that change could have a non-oscillatory temporal profile (Figure 7a). Of particular interest would be a ramp-to-step profile in which aspects of the middle or inner ear shift from one state to another during the course of a saccade and hold steady at the new state during the subsequent fixation period. This kind of temporal profile would match the time course of the saccade itself.

**Figure 7.**
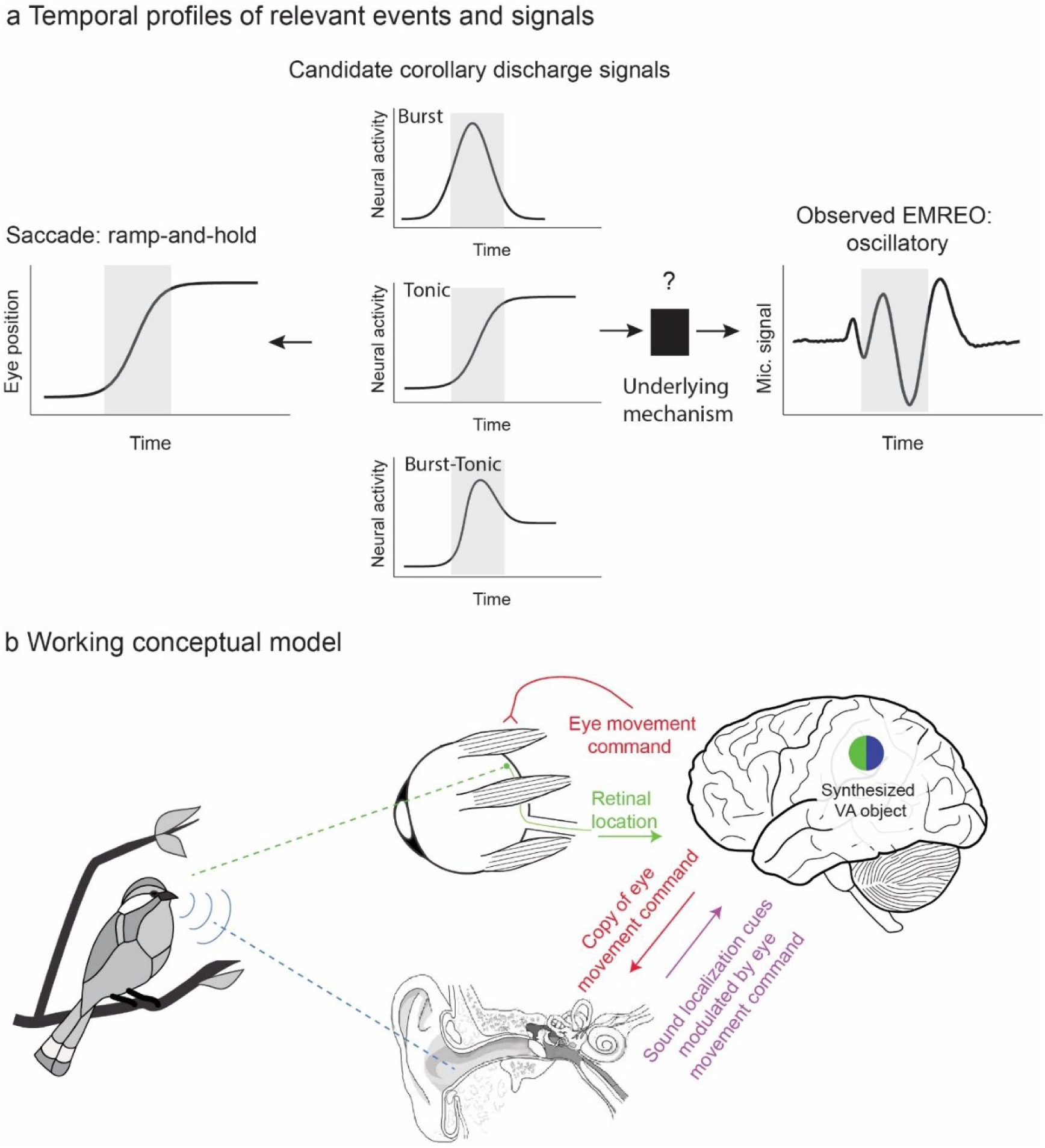
Temporal profiles of relevant signals and working conceptual model for how EMREOs might relate to our ability to link visual and auditory stimuli in space. **a.** Temporal profiles of signals. The EMREO is oscillatory whereas the eye movement to which it is synchronized involves a ramp-and-hold temporal profile. Candidate source neural signals in the brain might exhibit a ramp-and-hold (tonic) pattern, suggesting a ramp-and-hold-like underlying effect on an as-yet-unknown peripheral mechanism, or could derive from other known temporal profiles including bursts of activity time-locked to saccades. **b.** Working conceptual model. The brain causes the eyes to move by sending a command to the eye muscles. Each eye movement shifts the location of visual stimuli on the retinal surface. A copy, possibly a highly transformed one, of this eye movement command is sent to the ear, altering ear mechanics in some unknown way. When a sound occurs, the ascending signal to the brain will depend on the combination of its location in head-centered space (based on the physical values of binaural timing and level differences and spectral cues) and aspects of recent eye movements and fixation position. This hybrid signal could then be read-out by the brain.

Available eye movement control signals in the oculomotor system include those that follow this ramp-and-hold temporal profile, or tonic activity that is proportional to eye position throughout periods of both movement and fixation. In addition to such tonic signals, oculomotor areas also contain neurons that exhibit burst patterns, or elevated discharge in association with the saccade itself, as well as combinations of burst and tonic patterns (for reviews, see 59, 60). It remains to be seen which of these signals or signal combinations might be sent to the auditory periphery and where they might come from. The paramedian pontine reticular formation (PPRF) is a strong candidate for a source, having been implicated in providing corollary discharge signals of eye movements in visual experiments (61) (see also 62), and containing each of these basic temporal signal profiles (59, 60). Regardless of the source and nature of the descending corollary discharge signal, the oscillations observed here should be thought of as possibly constituting a biomarker for an underlying, currently unknown, mechanism, rather than necessarily the effect itself.

Despite these critical unknowns, it is useful to articulate a working conceptual model of how EMREOs might facilitate visual and auditory integration (Figure 7b). The general notion is that, by sending a copy of each eye movement command to the motor elements of the auditory periphery, the brain keeps the ear informed about the current orientation of the eyes. If, as noted above, these descending oculomotor signals cause a ramp-to-step change in the state of tension of components within the EMREO pathway, time-locked to the eye movement and lasting for the duration of each fixation period, they would effectively change the transduction mechanism in an eye position/eye movement dependent fashion. In turn, these changes could affect the latency, gain, or frequency-filtering properties of the response to sound. Indeed, intriguing findings from Puria and colleagues (35) have recently indicated that the tension applied by the middle ear muscles likely affects all three of these aspects of sound transmission throughout the middle ear. In short, the signal sent to the brain in response to an incoming sound could ultimately reflect a mixture of the physical cues related to the location of the sound itself - the ITD/ILD/spectral cues - and eye position/movement information.

Most neurophysiological studies report signals consistent with a hybrid code in which information about sound location is blended in a complex fashion with information about eye position and movement, both within and across neurons (6, 10, 11, 18, 19, 21, 44). Computational modeling confirms that, in principle, these complex signals can be “read out” to produce a signal of sound location with respect to the eyes (10). However, substantive differences do remain between the observations here and such neural studies, chiefly in that the neural investigations have focused primarily on periods of steady fixation. A more complete characterization of neural signals time-locked to saccades is therefore needed (8, 63).

Note that this working model differs from a spatial attention mechanism in which the brain might direct the ears to “listen” selectively to a particular location in space. Rather, under our working model, the response to sounds from any location would be impacted by peripheral eye movement/position dependence in a consistent fashion across all sound locations. However, such a system could well work in concert with top-down attention, which has previously been shown to impact outer hair cells even when participants are required to fixate and not make eye movements (64–70).

Another question concerns whether EMREOs might actually impair sound localization, specifically for brief sounds presented during an eye movement. We think the answer to this is no. Boucher et al (2) reported that perisaccadic sound localization is quite accurate, which suggests that EMREOs (or their underlying mechanism) do not impair perception. This is an important insight because given the rate at which eye movements occur - about 3/sec – and with each associated EMREO signal lasting 100 ms or longer (due to extending past the end of saccades, as explored by Gruters, Murphy et al. 2018 (25) and (28)), it would be highly problematic if sounds could not be accurately detected or localized when they occur in conjunction with saccades. If there is indeed a step-ramp system underlying the observed oscillations, then transduction of all sounds will be affected, regardless of when they occur with respect to saccades. Indeed, recent work supports the view that sound detection is unaffected by saccades (27).

All this being said, a role for EMREOs in computing the spatial location of sounds with respect to the visual scene does not preclude other roles. Specifically, they could also play a role in synchronizing sampling in the temporal domain (e.g. (29–33)). Such a possibility could account for the significant constant term C(t) of the regression analysis (Equation 1, Figure 3). This temporally precise non-spatial component could play a role in resetting or refreshing of auditory processing across time or coordinating with the refreshing of the visual image on the retina.

Overall, how brain-controlled mechanisms adjust the signaling properties of peripheral sensory structures is critical for understanding sensory processing as a whole. Auditory signals are known to adjust the sensitivity of the visual system via sound-triggered pupil dilation (71), indicating that communication between these two senses is likely to be a two-way street. The functional impact of such communication at low-level stages is yet to be fully explored and may have implications for how individuals compensate when the information from one sensory system is inadequate, either due to natural situations such as noisy sound environments or occluded visual ones, or due to physiological impairments in one or more sensory systems.

## Acknowledgments

We are grateful to Dr. Matthew Cooper, Dr. Kurtis Gruters, Jesse Herche, Dr. David Kaylie, Dr. Jeff Mohl, Dr. Shawn Willett, Meredith Schmehl, Dr. Jonathan Siegel, Chadbourne Smith, Dr. David Smith, Justine Shih, Chloe Weiser, Tingan Zhu, for discussions and other assistance concerning this project. This work was supported by NIH (NIDCD) grant DC017532 to JMG.

## Supplementary Materials

### Table of Contents

- Methods
- Supplementary Table – summary of included data
- Supplementary Figure 1
- Supplementary Figure 2
- Supplementary Figure 3 – supplement to Figure 1
- Supplementary Figure 4 – supplement to Figure 2
- Supplementary Figure 5 – supplement to Figure 4
- Supplementary Figure 6 – supplement to Figure 5
- Supplementary Figure 7 – supplement to Figure 6

## Methods

### General

Healthy human subjects that were 18 years of age or older with no known hearing deficits or visual impairments beyond corrected vision were recruited from the surrounding campus community (N=16; 8 female, 8 male; N=10 subjects tested for each task as described in the next section; female-male ratio was also equal in these subgroups). If subjects were unable to perform the saccade task without vision correction, they were excluded from the study. All study procedures involving subjects were approved by the Duke Institutional Review Board, and all subjects received monetary compensation for their participation.

Acoustic signals in both ear canals were measured simultaneously with Etymotic ER10B+ microphone systems coupled with ER2 earphones (Etymotic Research, Elk Grove Village, IL) to allow calibrations of the microphones (however, note that no auditory stimuli were used during any of the saccade tasks in the current study). A low-latency audio interface (Focusrite Scarlett 2i2, Focusrite Audio Engineering Ltd., High Wycombe, UK) was used for audio capture and playback through the Etymotic hardware at a sampling rate of 48kHz. Eye tracking was performed with an Eyelink 1000 system sampling at 1000Hz. Stimulus presentation and data acquisition were controlled through custom scripts and elements of The Psychophysics Toolbox in MATLAB, with visual stimuli presented on a large LED monitor.

In all experiments, eye position and microphone data were recorded while subjects performed silent, visually-guided saccade tasks. Experimental sessions were carried out in a darkened, acoustically isolated chamber made anechoic with the addition of acoustic damping wall tiles. Subjects were seated 70 cm from the screen, and a chin rest was used to maintain head position and minimize movement. Experimental sessions were subdivided into multiple runs, approximately 5 minutes each. This provided subjects with the opportunity to take a brief break from the experiment if needed to maintain alertness or to address any possible discomfort from maintaining their posture. Each run typically consisted of approximately 125 trials and fixation positions and saccade targets were presented in pseudorandom order.

Before each experimental session, the eye-tracking system was calibrated using the calibration routine provided with the Eyelink system to register raw eye-tracking data to gaze locations on the stimulus presentation screen. If the subject requested an adjustment to the chin rest or left the recording chamber for a break, the eye-tracking calibration was repeated. Before each run, the microphone system was calibrated to ensure that each microphone had a frequency response that was similar to the pre-recorded frequency response of the microphones when placed in a volume that approximated the size and geometry of the human ear canal - a 3ml syringe cut to accept the Etymotic earpieces. The syringe stopper was pulled to 1.25 cm^3^ to approximate the volume of the average adult human ear canal. A small amount of gauze (.25cm^3^) was added to the volume to emulate the attenuation caused by the soft tissue of the ear canal. The calibration routine played tones from 10 to 1600Hz, at a constant system output amplitude. As the purpose of this calibration was to compare microphone function in a control volume with that in an earpiece just placed in a subject, the weighting of the tones was not specifically calibrated. If the input-output results of the same tone sequences were consistent between ears and matched the overall shape of the syringe calibration curves, microphone placement was considered successful. No sounds were delivered during periods of experimental data collection.

### Task descriptions

All tasks followed the same stimulus timing sequence: initial fixation points were displayed on screen for 750ms and then removed as the saccade targets were presented for 750ms (Supplementary Figure 1A). Fixation and target locations were indicated by green dots. Subjects were instructed to fixate on the initial fixation locations until targets were presented on the screen, then to saccade to the targets and fixate on the targets until they changed from green to red for the last 100ms of the target presentation (the color cue was intended to help subjects maintain fixation through the end of the target presentation). Inter-trial-intervals were jittered 350±150ms. This was done to minimize the potential impact of non-saccade related noise signals that may be periodic (i.e. heartbeat, external acoustic and electromagnetic sources).

In the five-origin grid task (Supplementary Figure 1B), participants performed saccades to multiple targets from five different initial eye positions in a plus-shaped configuration at −9°, 0°, and +9° horizontally and at −6 °, 0°, and 6 ° of elevation as shown. Twenty five saccade targets ranged from −18° to +18° in 9° degree increments horizontally and from −12° to +12 ° in 6 ° increments vertically.

In the horizontal/vertical task (Supplementary Figure 1D), participants performed saccades to targets along the vertical and horizontal axes from a central fixation. Vertical targets ranged from −12° to +12° in 3° increments and horizontal targets ranged from −18° to +18° in 3° increments.

In the single-origin grid task (Supplementary Figure 1C), participants made saccades to 24 distinct targets of varying vertical and horizontal placement combinations from a central fixation. Horizontal location components ranged from −18° to +18° in 9° increments and vertical location components ranged from −12° to +12° in 6° increments.

### Preprocessing analysis

#### Saccade-microphone synchronization

Microphone data was synchronized to the onset the saccade from the fixation point to the target. This was defined based on the third derivative of eye position, or jerk. The first peak in the jerk represents the moment when the change in the eye acceleration is greatest. Prior to each differentiation, a lowpass discrete filter with a 7ms window was used to smooth the data and reduce the effects of noise and quantization error. This filter was normalized, such that its output to a constant series of values equaled those values.

#### Trial exclusion criteria

Trials were excluded based on saccade performance and the quality of microphone recordings. Exclusion criteria used for eye tracking: 1) if subjects made a sequence of two or more saccades to achieve the target; 2) if the eye tracking signal dropped out during the trial (e.g. due to blinks); 3) if the eye movement was slow and drifting, rather than a saccade; 4) if the saccade curved by more than 4.5° (subtended angle); or 5) subjects failed to maintain 200ms of fixation before and after the saccade. 6) If eye tracking dropped samples that prevented the calculation of the saccade onset time. On average these saccade-related exclusion criteria resulted in the exclusion of ∼15% of the trials.

Prior to any further analysis, microphone data was downsampled from 48 kHz to 2 kHz sampling rate to reduce processing time given that the previously observed eye-movement related signals of interest are well below 100 Hz (25). Exclusion based on noise in the microphone recordings was minimal. Within each block of trials, the mean and standard deviation of the RMS values for each trial was calculated. Individual trials were excluded if the microphone signal on that trial contained any individual values that were more than 10 standard deviations away from that mean. This typically resulted in the exclusion of a further ∼1.4% of the trials, after application of the eye position screen described above. On average, about 1200 trials were included for each subject and task condition. See Supplementary Table for a full breakdown across subjects and tasks.

### Z scoring

To facilitate comparison across subjects, sessions, and experiments, all microphone data reported in this study was z-scored within blocks and prior to the application of the exclusion criteria described above. The mean and standard deviation of the microphone values in a window −150 to −120 ms prior to saccade onset were used as the normalization baseline period.

### EMREO intensity estimates

In addition to the Z scoring, we also calculated the peak-equivalent sound pressure levels associated with EMREOs. EMREOs occur in a frequency range (∼40 Hz) for which the gain of the ER10B+ recording microphone is not flat, so standard conversions from voltage-to-pressure do not apply. Rather, we used a Bruel & Kjaer sound level meter (Model 2245) which has a roughly flat frequency response 20 Hz to 20 kHz (Z-weighting). Using this meter, we measured the sound pressure levels associated with 40 Hz pure tones generated by Etymotic ER-2 transducers coupled in the earbud with the ER10B+ microphone, which recorded these tones simultaneously. These measurements were made in a short length of tygon tubing simulating the approximate volume and sound-absorption properties of the human ear. This procedure provided a voltage-to-sound-pressure-level mapping in the 40 Hz frequency range. We then used these values to estimate the peak-equivalent sound pressure level of the EMREO signal for selected targets in individual subjects (Supplementary Figure 2). By this technique, EMREOs ranged in intensity from ∼49 to 64 dB SPL, with a mean of 56 dB SPL. These values closely corresponding with estimates using either the same (28) or a different method (25).

### Regression Analyses

Regression was used to assess how EMREOs vary with both eye position and eye movement. Linear methods were used as a conservative approach providing a lower bound on the ability to account for the variation in the microphone signal on the basis of eye movement parameters.

The microphone signal at each moment in time *Mic(t)* was fit as follows:

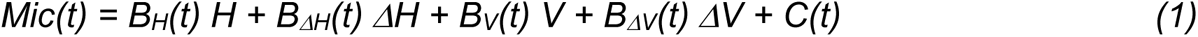

where *H* and *V* correspond to the initial horizontal and vertical eye position and *ΔH* and *ΔV* correspond to the respective changes in position associated with that trial. The slope coefficients *B_H_, B_ΔH_, B_V_, B_ΔV_*are time-varying and reflect the dependence of the microphone signal on the respective eye movement parameters. The term *C(t)* contributes a time-varying “constant” independent of eye movement metrics, and can be thought of as the best fitting average oscillation across all eye positions and displacements.

The term *C(t)* was included for all regressions, but other parameters were omitted when not relevant. Specifically, for the single-origin grid tasks and horizontal-vertical tasks, the model used vertical and horizontal saccade displacement (*B_ΔH_(t) ΔH*, *B_ΔV_(t) ΔV)* as regression variables but not *B_H_(t) H* or *B_V_(t) V* as initial position did not vary for those tasks. The analysis produced values for the intercept and variable weights (or slopes), their 95% confidence intervals, R^2^, and p-value for each time point.

For most analyses, the measured eye positions/changes-in-eye positions were used as the independent variables, so as to incorporate any variability due to scatter in fixation or saccade endpoint. For the target readout analysis described in Figures 5 and 6, the horizontal and vertical positions of the targets, rather than the associated eye movements, were used.

## Data Availability

The data and analysis code for this paper will be made available on publication on Duke University’s Research Data Repository. (citation to be added).

---

**Supplementary Table.**
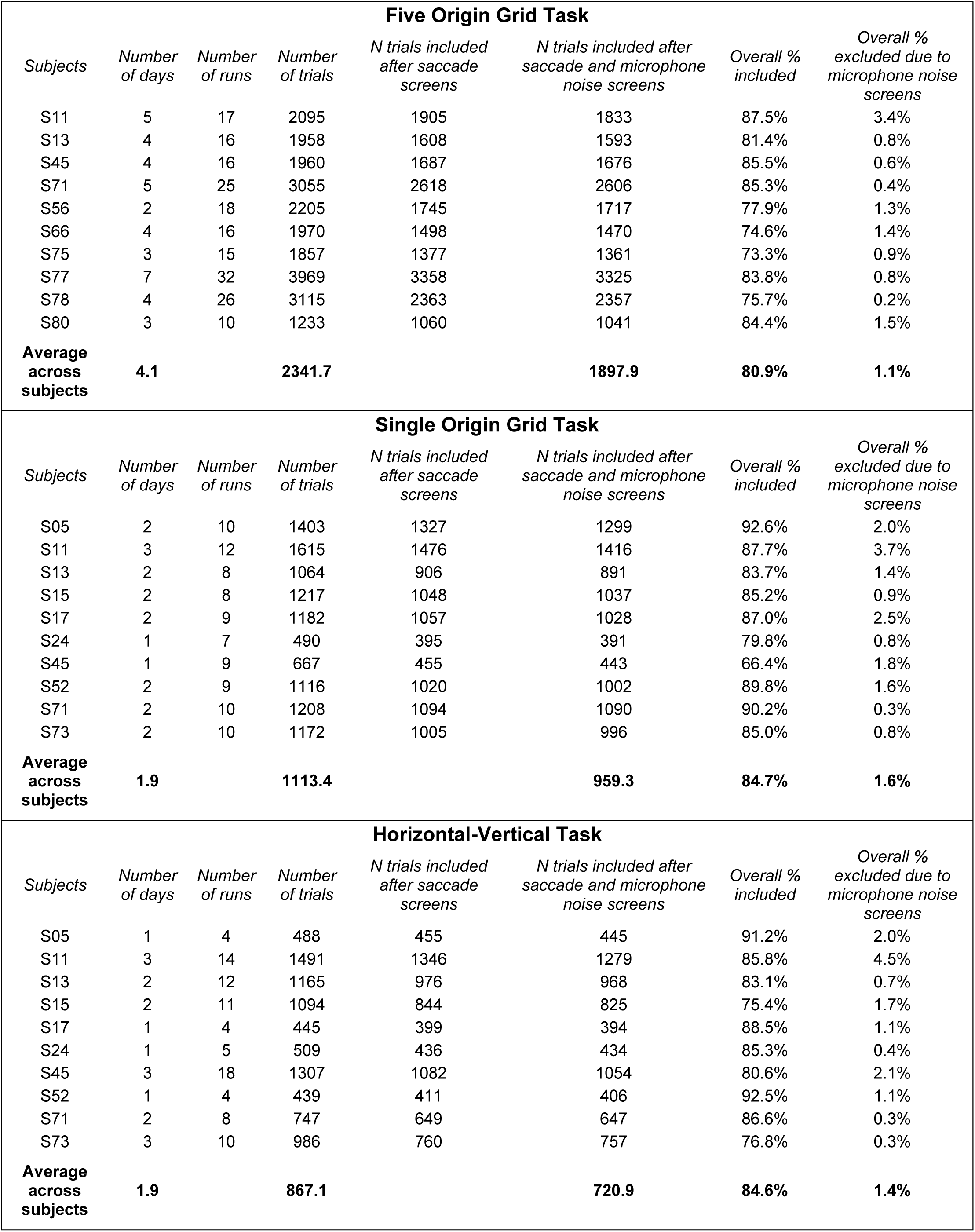
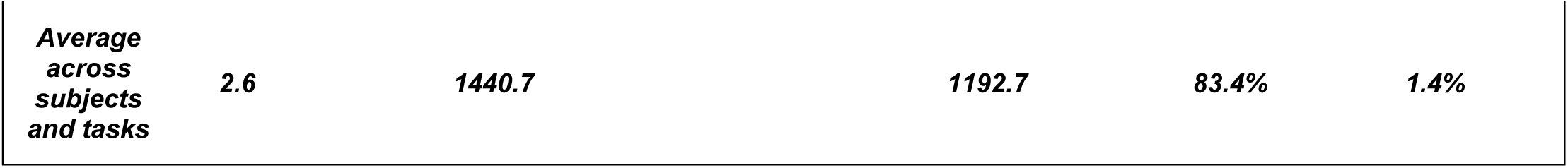
The numbers of days, blocks, and trials recorded and included/excluded based on saccade parameters and/or microphone noise across the different tasks.

**Supplementary Figure 1.**
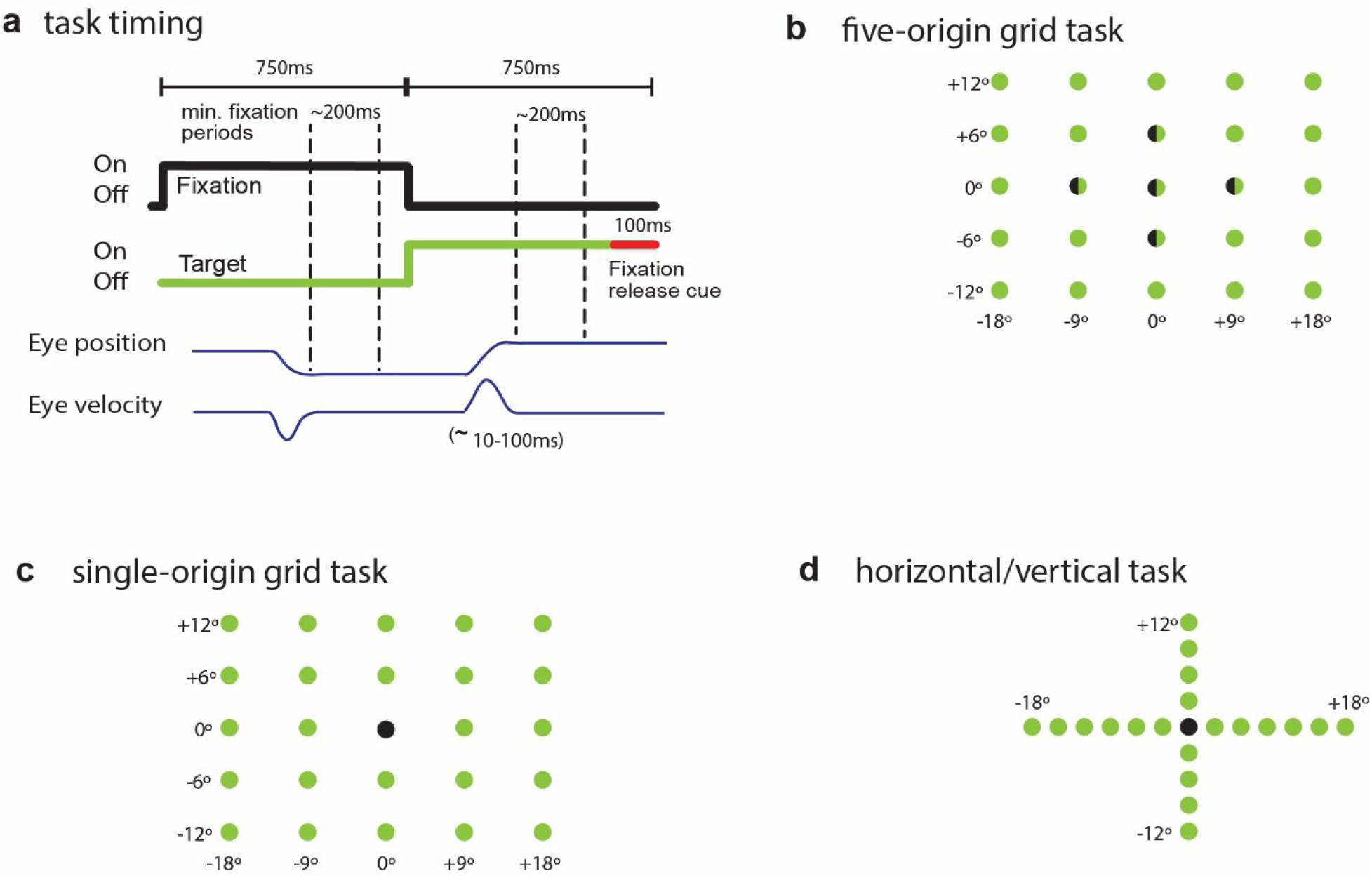
Events of the tasks in time and space. a. Task events across time. Each trial began with the onset of an initial “<u>fixation</u>” cue (black trace). Participants made saccades to the fixation point, then maintained fixation for a minimum of 200 ms. The fixation point was then turned off and a new “<u>target</u>” was turned on (green trace). Participants saccaded to this target and fixated for another 200 ms, at which point the target turned red indicating that the trial was over. The ear-canal recordings were analyzed in conjunction with the fixation-point-to-target saccade. b-d. Spatial layouts of fixation points and targets for the three task designs used in this study. Points in space that were used as both a fixation and a target across different trials are half-green, half-black.

**Supplementary Figure 2.**
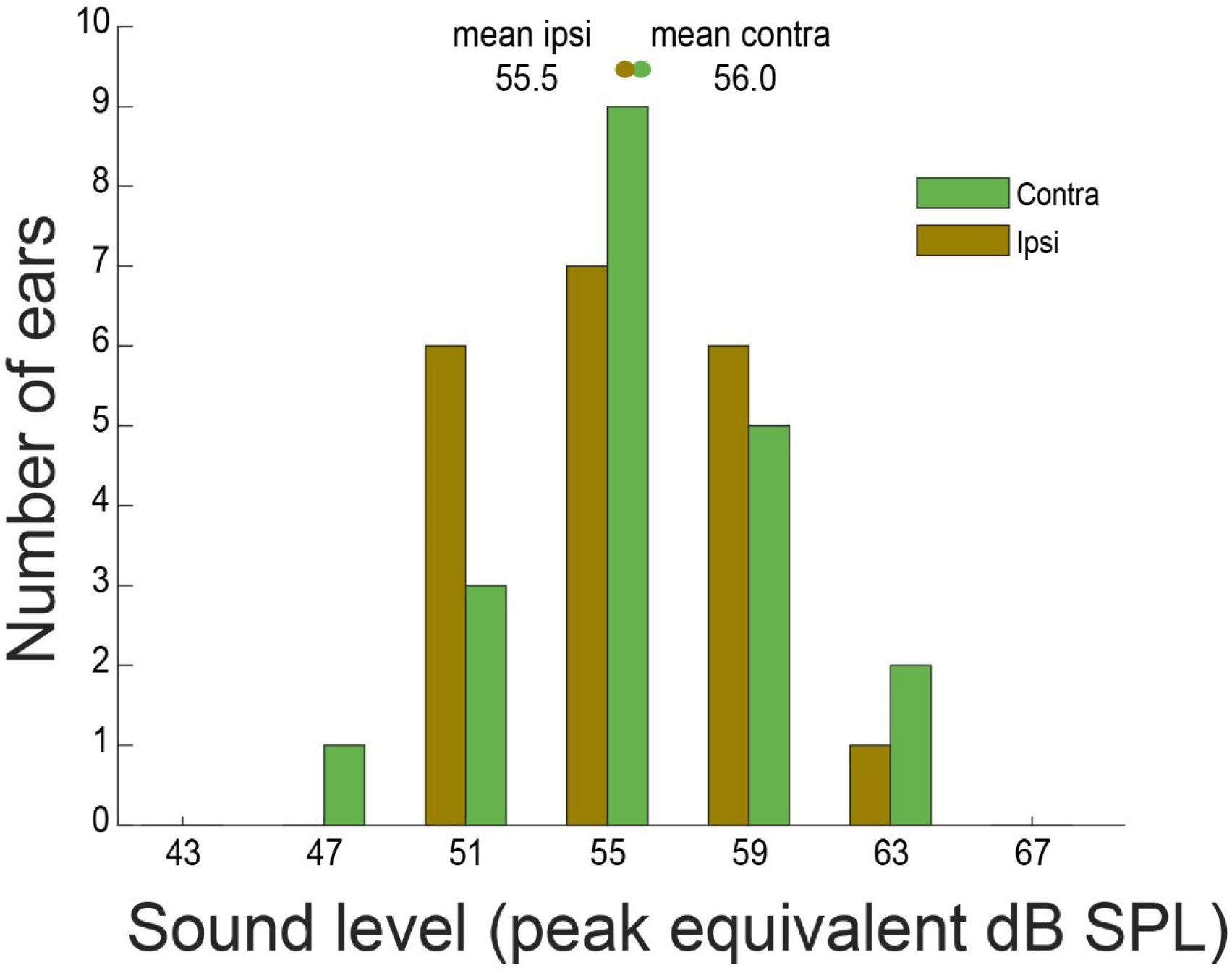
Sound level of EMREOs associated with the 18 degree contralateral or ipsilateral target locations for individual subjects tested in the five-origin grid task. The origin for these data was the (0, −6) origin which corresponds to the conditions tested in our earlier study (25). See “Methods: EMREO intensity estimates” for details.

**Supplementary Figure 3.**
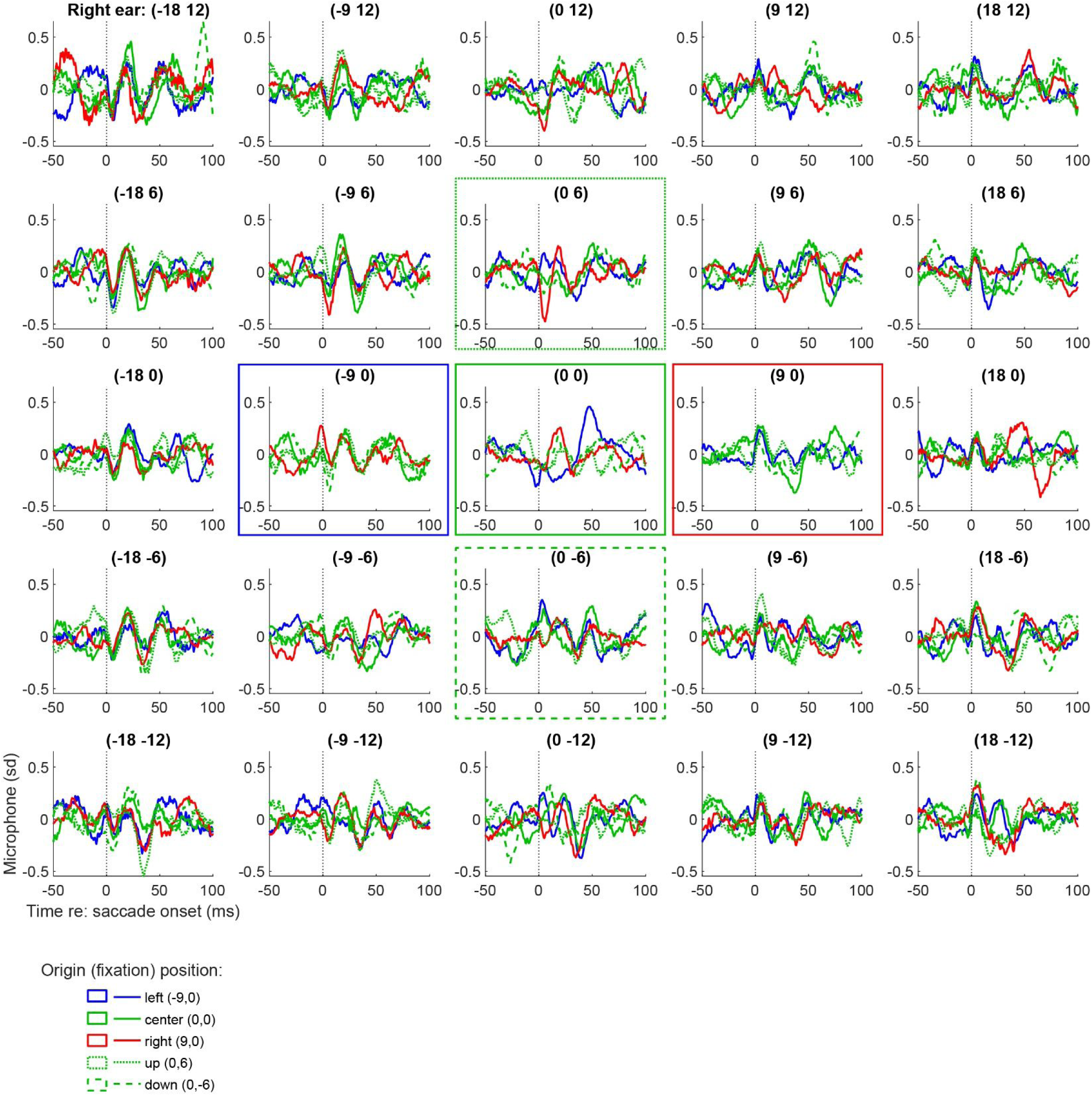
Grand average EMREOs recorded during the 5-origin grid task in right ears of ten subjects. Same format as Figure 1: Each panel is the grand average EMREO signal (average of the individual subject averages) that is generated when a saccade is made to that location on the screen e.g. the top right panel involves saccades to the top right target location. The color and line styles of each trace correspond to the initial fixation point as indicated by boxes of the same color and line style; e.g. all red oscillations are generated during a simultaneous saccade that originated from the right fixation point.

**Supplementary Figure 4.**
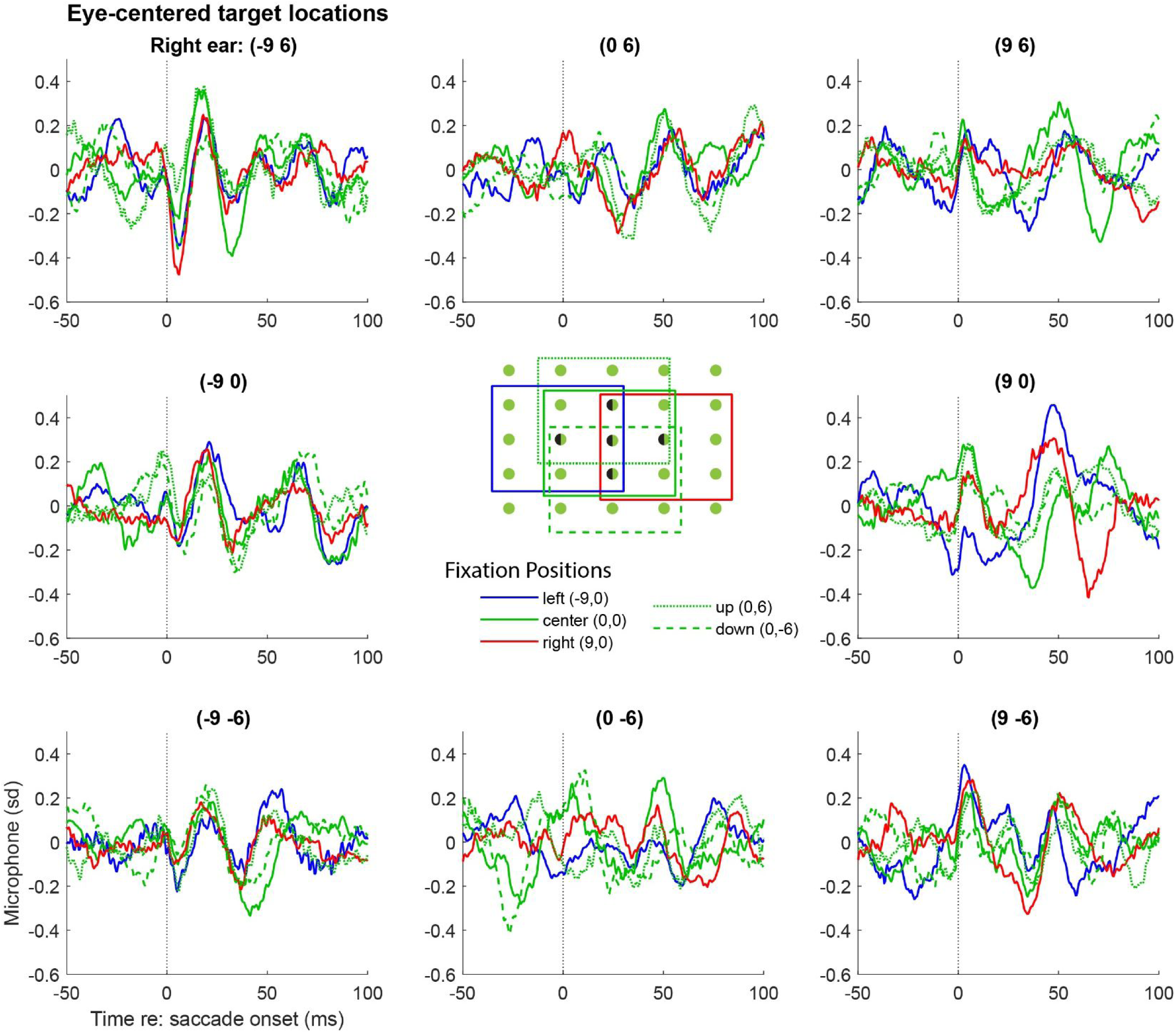
Grand average EMREOs as a function of target location with respect to the fixation position, for N=10 right ears. Same format as Figure 2. The data shown are the same as (a subset of) those shown in Supplementary Figure 3, but here each panel location corresponds to a particular target location defined with respect to the fixation point. The color/linestyle indicate the associated fixation position, as in Supplementary Figure 3.

**Supplementary Figure 5.**
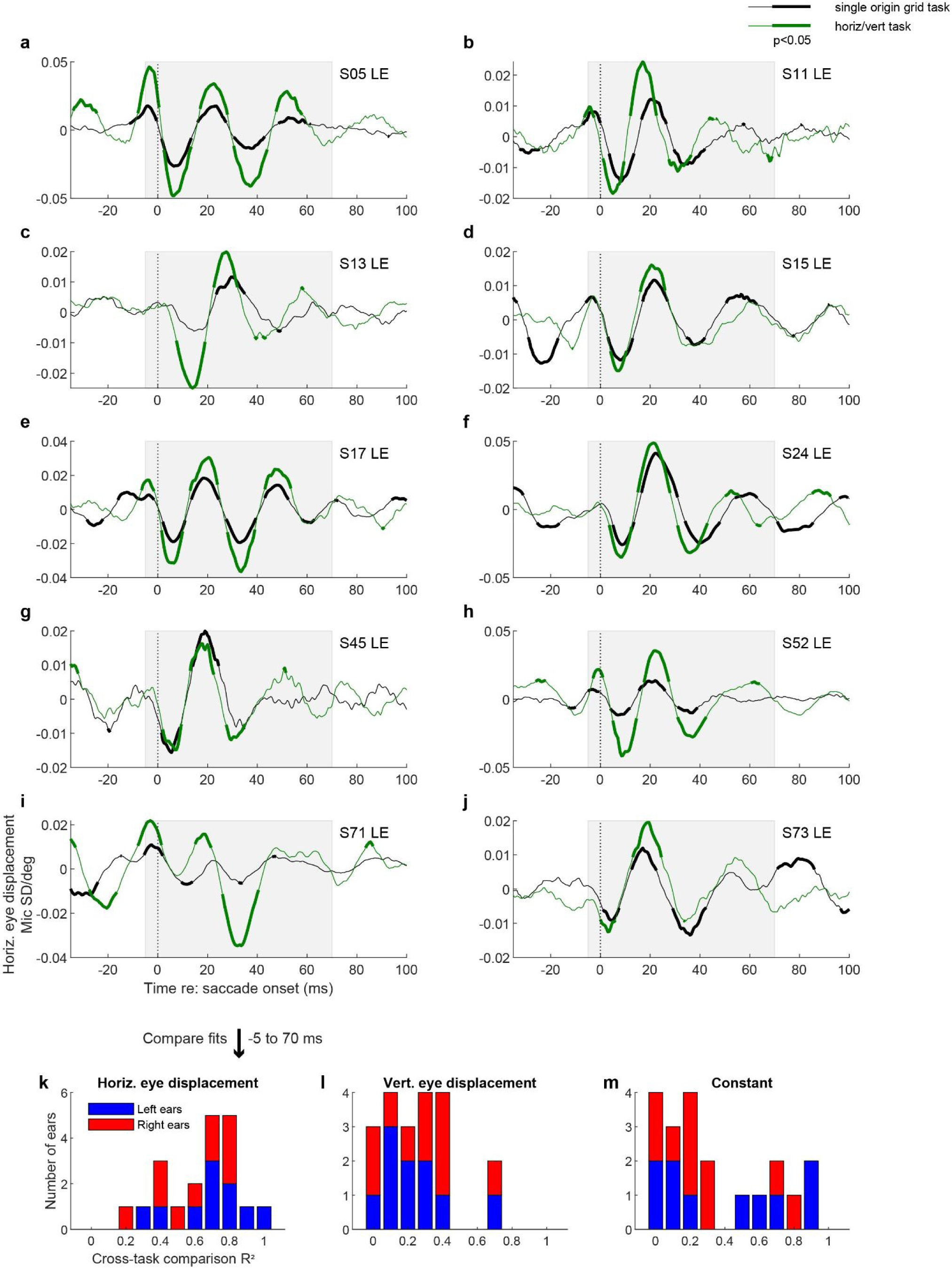
Cross-task comparison for individual subjects and at the population level. A-J. Regression coefficients for horizontal eye displacement in the single-origin grid task (black) vs. the horizontal-vertical task (green) for individual subject left ears, similar to Figure 3A. K-L. All three coefficient values (horizontal eye displacement, vertical eye displacement, and constant terms) were then (separately) regressed against each other for the two tasks during the time window −5 to 70 ms (grey boxes in panels A-J), and the distribution of R^2^ values across subjects and ears are shown as indicated. There is excellent cross task correspondence for the horizontal eye displacement signal at the individual level and somewhat more variable correspondence for the vertical eye displacement and constant terms, consistent with the population results shown in Figure 4.

**Supplementary Figure 6.**
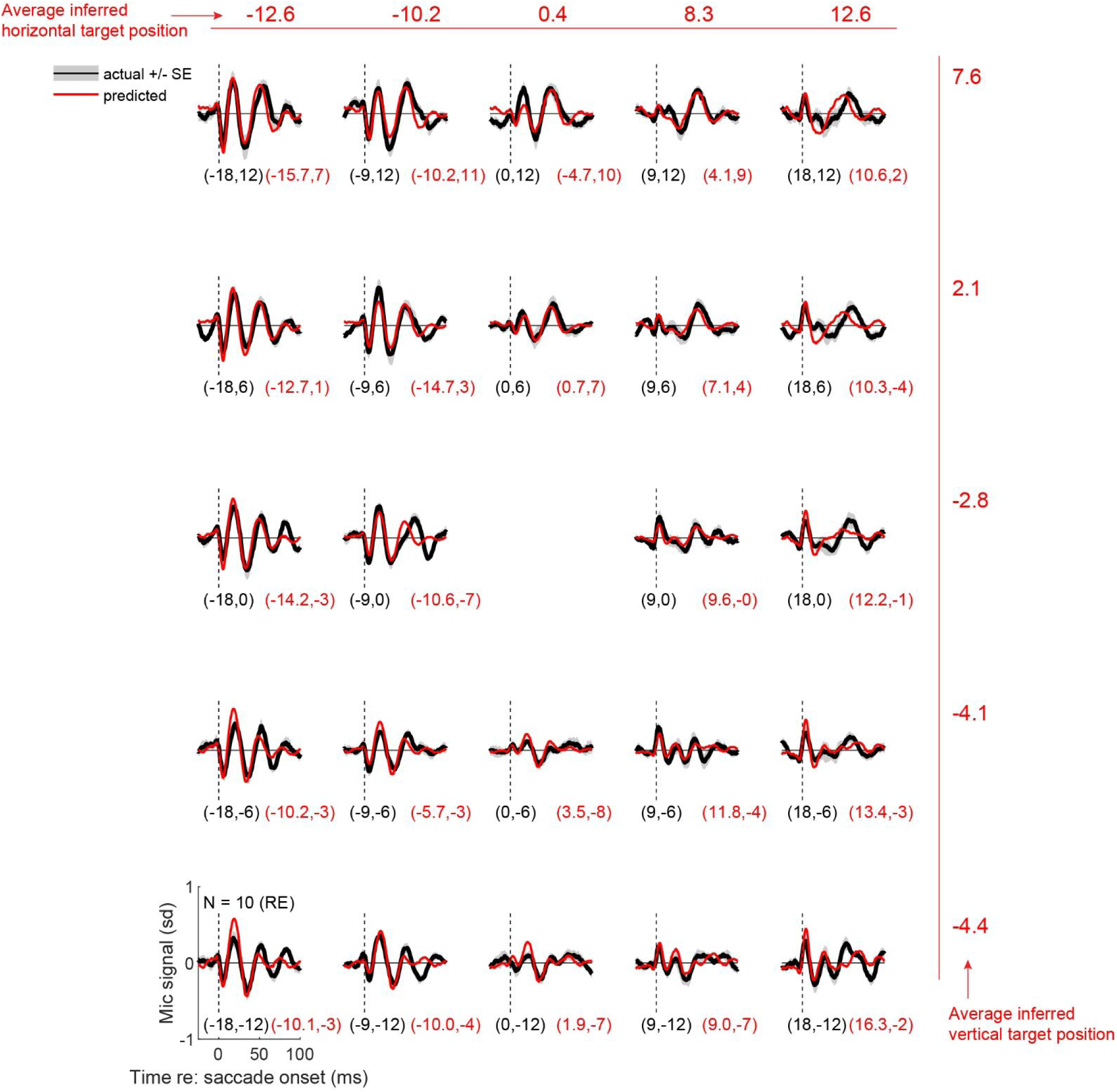
Same as Figure 5, but for right ear data. See main text for details.

**Supplementary Figure 7.**
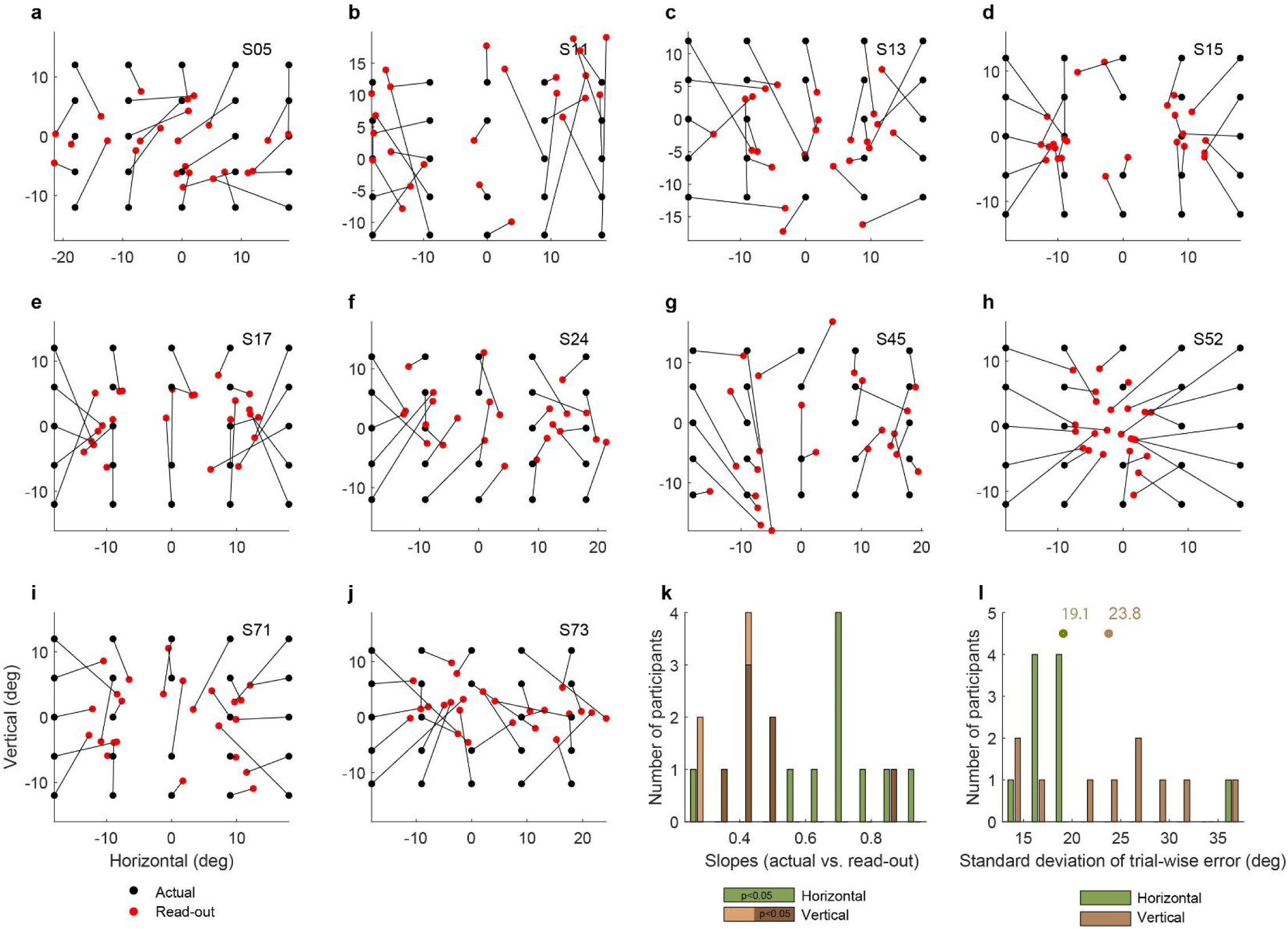
Target readout results for individual subjects following the population results shown in Figure 6J. The readout was computed based on the difference between the two ears for the horizontal dimension and the average between the two ears for the vertical dimension. Panels A-J show the results for each subject. Panel K shows the distribution of slope values relating the actual vs. read-out target locations in the horizontal and vertical dimensions (green: horizontal; brown: vertical). All values were statistically significant p<0.05 in the horizontal dimension and most were significant in the vertical dimension (darker shade of brown). The horizontal values are comparable to the population values indicated in Figure 6H, and the vertical are comparable to Figure 6K. Panel L shows the average standard deviations of the difference between the read-out location and the actual target when the readout is computed from individual trial waveforms.

## Footnotes

1 We note that EMREOs are unlikely to be due to the actual sound of the eyes moving in the orbits. Our original study, Gruters et al (2018) showed that when microphone recordings are aligned on saccade offset (as opposed to onset, as we did here), EMREOs continue for at least several 10’s of ms after the eyes have stopped moving.

## Notes

### Competing Interest Statement

The authors have declared no competing interest.

### Summary of Updates

Figure 1 moved to the Supplementary Figures; subsequent figures unchanged except for the numbering. References updated

